# Construction of mutant heparinase I with significantly increased specific activity

**DOI:** 10.1101/2020.05.12.092361

**Authors:** A.N. Kalinina, L.N. Borschevskaya, T.L. Gordeeva, E. Patrusheva, S.P. Sineokiy

## Abstract

The cleavage of heparin by heparin lyases showed great potential as a cost-effective and innoxious method for producing heparin with low molecular weight (LMWH). One of the most studied and sought heparin lyase is heparinase I (HepI). However, the industrial use of HepI was largely hampered by its low specific activity and thermal stability. In this article we describe increasing in specific heparinase I activity by stepwise site-directed mutagenesis. Thus after two cycles of mutagenesis, we obtained mutant heparinase I Flavobacterium heparinum with significantly increased specific activity (25%).

---

Heparin, a sulfated glycosaminoglycan, - a general name of a heterogeneous mixture of sulfated polysaccharide chains, which include repeating units of D-glucosamine, L-iduronic or glucuronic acid (Yang et.al.1985).

Heparin is used as a critical anticoagulant in clinical practice (Casu and Lindahl, 2001; Lever and Page, 2002). It should be noted that only about 30% of the molecules have heparin anticoagulant activity. Low molecular weight heparin (LMWH) has better pharmacokinetics, bioavailability and safety compared to unfractionated heparin. LMWH is obtained by chemical or enzymatic degradation of heparin (Linhardt and Gunay, 1999). A promising proposal in most thromboembolic treatments is the replacement of unfractionated heparin with LMWH (Hirsh et al., 2001; Sundaram et al., 2003).

Treatment of heparin by heparinase is the most promising way to obtain LMWH.

Heparin lyases (heparinase) are enzymes which cleave heparin and heparan sulfate on lycosidic linkages between hexosamine and uronic acids via a b-elimination mechanism (Ernst et al., 1995). Heparinases were first isolated and characterized from Pedobacter heparinus (formerly Flavobacterium heparinum) (Yang et al., 1985),

It should be noted, that the most rational LMWH - drugs with a molecular weight of 4000-7000 Daltons. Such mucopolysaccharides are obtained by treatment of heparinase I on heparin (Wallentin, 1996; Hollingsworth et al., 2000; Dunn et al., 1996).

Heparinase I (Hep I), one of the most widely studied glycosaminoglycan lyases, specifically cleaves heparin on the links between D-glucosamine and sulfated L-iduronic acid (Ernst et.al. 1995)

However, the use of HepI has not received industrial use because of its low specific activity and stability. (Korir and Larive, 2009; Sasisekharan et al., 1993; Shpigel et al., 1999).

For the first time the successful realize the soluble expression of HepI in E. coli, was carried by fusion HepI to maltose-binding protein (MBP) and succeeded in its high-yield production and easy purification. As a result, approximately 90% of protein was in soluble heparinase active form. And in this case protein had specific activity of 88.3 units / mg of protein (reducing specific activity by 25%) (Chen et al., 2005, 2007).

Nevertheless, even in the form fusion of maltose binding protein with heparinase I enzyme shows low thermostability and specific activity.

The C297S mutation had been introduced in the gene encoding heparinase I, in order to increase the thermostability of the enzyme MBP-HepI, as well as the reaction conditions were chosen the degradation of heparin. In the course of this work showed a significant increase in the thermostability and residual enzymatic activity after prolonged storage (Chen et al.,2011, 2013).

Our group is involved in work the cost-effective and industry-applicable catalysis with heparinase I through improving its productivity and activity. The purpose of our work was increasing the specific activity of the enzyme, based on the data structure of the active site and mechanism of action of heparinase I.

Amino acid residues cysteine - 135 and histidine-203 of heparinase I play a major role in the cleavage of heparin. Thus, during the initiation of the reaction, the sulfur of the thiol group on the cysteine-135 is formed a negative charge, then the thiolate anion performs nucleophilic attack heparin, during which are formed the breaking of bonds in the oligomers and form mukopolisaaharides. It should be noted that increasing the specific activity can be realized by improving in enzyme stability in the form of the cysteine-135 thiolate anion (Sasisekharan, 1991).

Our task was a search effective areas in sequence heparinase I and the introduction of mutations that could lead to the stabilization of the negative charge on the thiol group of cysteine - 135 in order to increase the specific activity of enzyme.

As a result was obtained the mutant enzyme heparinase I with a significantly increased specific activity (approximately 25%).

## Materials and methods

### Materials, bacterial strains, plasmids and media

Heparin sodium salt (150 I.U./mg) was purchased from ApplliChem. Heparinase I from *Flavobacterium heparinum* (≥200 U/mg) was purchased from Sigma-Aldrich. F. heparinum (ATCC 13125) was purchased from the Institute of Applied Microorganisms (IAM) of the University of Tokyo, Japan. E. coli Tuner DE3 was purchased from VKPM (Russia). This strain was used for studying influence introduced mutations in a gene HepI on activity of the enzyme. Plasmid pET22b(+) preserved in our laboratory was used.

F. heparinum was grown at 30 ° C in B1 medium which contained 3g/l NaCl, 3g/l beef extract and 10g/l tryptone. E. coli Tuner DE3 was grown in Luria Bertani (LB) media with or without ampicillin (100 μg/ml).

### Construction of the expression vector for native and mutant heparinase I

Genomic and plasmid DNA isolation, restriction and ligation of DNA fragments and transformation followed the standard procedures of Molecular Cloning (Sambrook J. and Russel D.W. 2001).

F. heparinum genome DNA was isolated using the Genomic DNA Purification Kit (Fermentas). Plasmid was prepared by GeneJET Plasmid Miniprep Kit (Fermentas). Heparinase I (HepI) gene was amplified from the isolated genome DNA by the oligonucleotides listed in table 1.

**Table 1.**
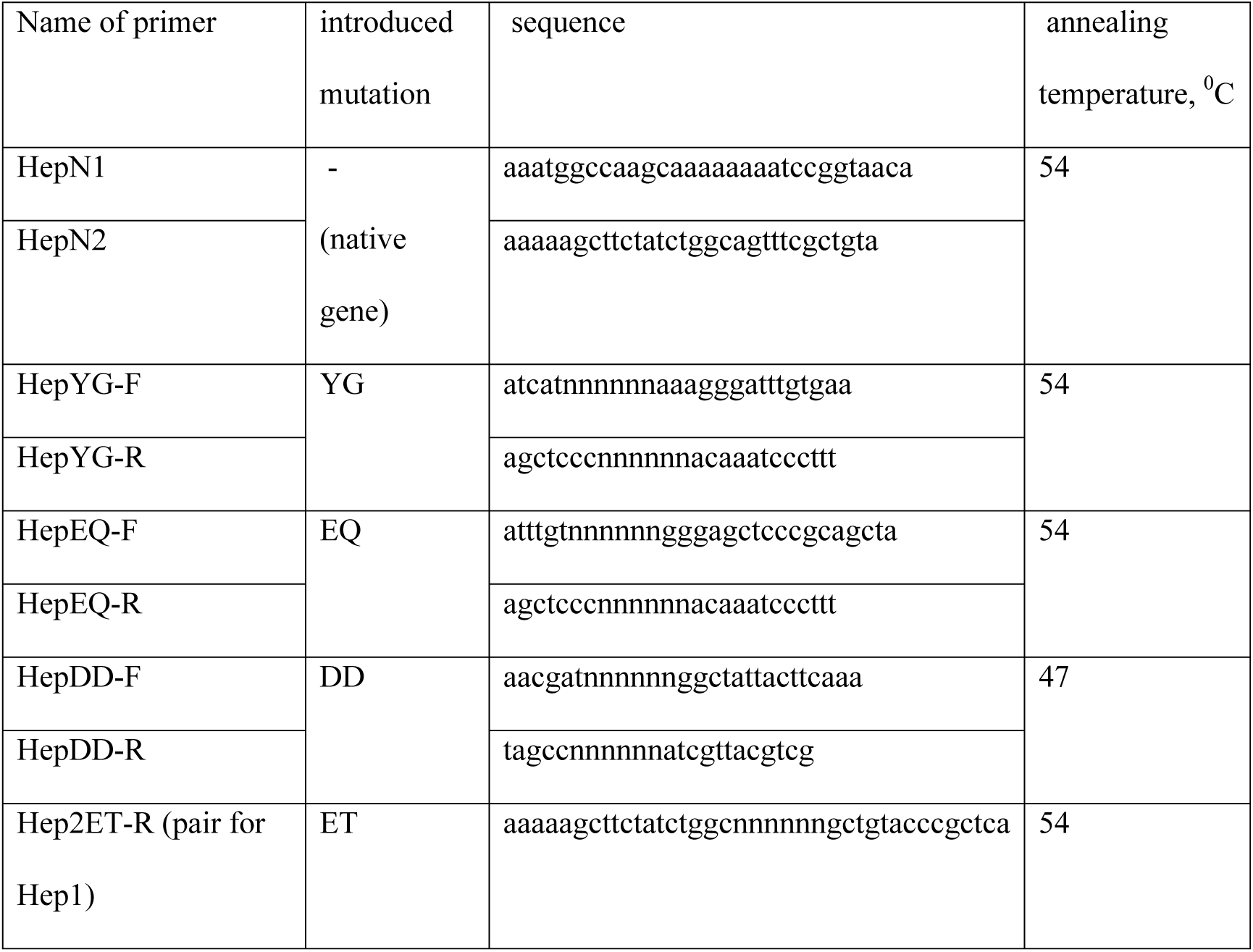
Primers for amplification and introduction mutations gene Hep I

To construct pET-HepN, the pair primers HepN1 and HepN2 were used to generate a HepI native gene with MlsI and HindIII restriction sites at the 5’ and 3’ sites. After purification, the fragment was digested by restrictases. Then it was ligated with pET22b(+) which was digested with the same two enzymes and recovered from agarose gel.

The mutations were introduced using the PCR method of Higuchi (Higuchi, 1990). This method involved the following steps. Two primary PCR reactions (12 cycles) were set up using the pET-HepN construct as a template and the following combinations of primers: HepN1 primer and 3’ mutant primer, HepN2 primer and 5’ mutant primer. Then two produced parts of gene were mixed and full mutant gene HepI was amplified using primers HepN1 and HepN2. After purification, the fragment was digested by restrictases MlsI and HindIII. Then also it was ligated with pET22b(+).

Constructions with fusion gene HepI with His6-Tag on the C-terminal end were obtained on base pET22b(+) plasmid for further purification of enzymes.

The recombinant plasmids of pET-Hep were transformed to E. coli Tuner DE3 to give E. coli Tuner DE3 (pET-Heps). Further, recombinant plasmids were isolated from the resulting strains, and mutant and native genes HepI were sequenced. Processing the received sequences were performed using the Vector NTI

### Functional expression of HepI

E. coli cells were cultured in a LB medium containing ampicillin (100 μ/ml). The cells were first grown at 37 °C in tubes (200 rpm) overnight. The 50 μl of overnight cultures were transferred into 5ml LB (without ampicillin) and grown 1,5 hour at 30 °C and 200 rpm. Then 100 μl 0,5 M isopropyl β-D -1-thiogalactopyranoside (IPTG) was added to culture and induced 2 hour at 30 °C and 200 rpm. Then the cells were harvested by centrifugation (1min at 4 °C and 5000 rpm). Total enzymatic activity assay

The total activity was measured for test a large number of mutant enzyme. The total heparinase I activity assay in supernatant was carried out according to the Azure A method, which was investigated in our group (see Results).

### Purification of HepI

To obtain values of specific activities purification of enzyme was conducted. The proteins were purified as described previously (Sasisekharan, R. et.al. 1995)

### Protein SDS–PAGE

Protein purity was confirmed on 12.5 % SDS-PAGE according to the general Laemmli method (Laemmli 1970) and system Mini Protean II (Bio Rad), for appearing of proteins were used Silver Stain Plus kit (Fermentas).

### Enzyme activity assay and protein determination

Heparinase activity was measured according to modification of the UV 232 nm method (Bernstein et al. 1988). The enzymatic reaction was carried out at 30 °C using heparin as the substrate in Tris–HCl buffer (pH 7.4) (containing 25 g/l heparin, 40 mM NaCl, 3.5 mM CaCl 2, and 17 mM Tris–HCl). Heparin degradation was monitored by following the UV absorbance at 232 nm on a Genesys 10S UV-VIS Therno scientific spectrophotometer and the activity was determined from calibration curve, which was constructed on the basis of data obtained by measurement of various concentrations of heparinase I (Sigma).

## Results and Discussion

The mechanism for the catalytic cleavage of heparin by heparinase I has been published previously. Histidine-203 during the initiation of catalysis activates cysteine-135 by separating the proton. Cysteine-135 acts as the base and separates the C5 proton from the urinate resulting in a 4-5 unsaturation. Then, histidine-203 protonates the leaving hexamine group. This completes the catalysis cycle. (Hirsh, J. et.al. 2001).

Previously, it was suggested that in order to activate the thiol group of cysteine 135 plays an important role the positive charge of the field environment by lowering its pKa. Thus, it can serve as the basis for proton abstraction (Sasisekharan et.al., 1995).

It was previously noted that heparinase I has similarities to the double specific and tyrosine-specific protein tyrosine phosphatase (PTPases) (Denu et.al., 1995). So, the highly reactive thiol group in the positively charged region is a common feature for these enzymes, and the role of the positively charged medium is not only in the binding of the substrate, but also in the activation of the thiol group for catalysis by reducing its pKa. In addition, both enzymes cleave a strongly negative charged substrate (Godavarti, 1996).,

Earlier, using the crystal structure of the active site Yersinia PTPase as an example, it was shown that thiol is stabilized as an anion by a complex network of hydrogen bonds with neighboring residues.

In order to reduce pKa for thiol, additional stabilization by surrounding positive residues is necessary (Zhang and Dixon, 1993).

Our work was focused on searching areas in environment of cysteine-135 thiol group heparinase I and making substitutions in them in order to improve stabilization of thiol group and increase heparinase I activity.

A three-dimensional model structure of P. heparinus HepI is unknown, but Protein data bank contain a three-dimensional model structure of *Bacteroides thetaiotaomicron* HepI. Sequence alignment was performed by ClustalW and showed that PhHepI shares 68% identity with the *B.thetaiotaomicron* HepI(Fig. 1) (Chenna et al., 2003).

**Figure 1.**
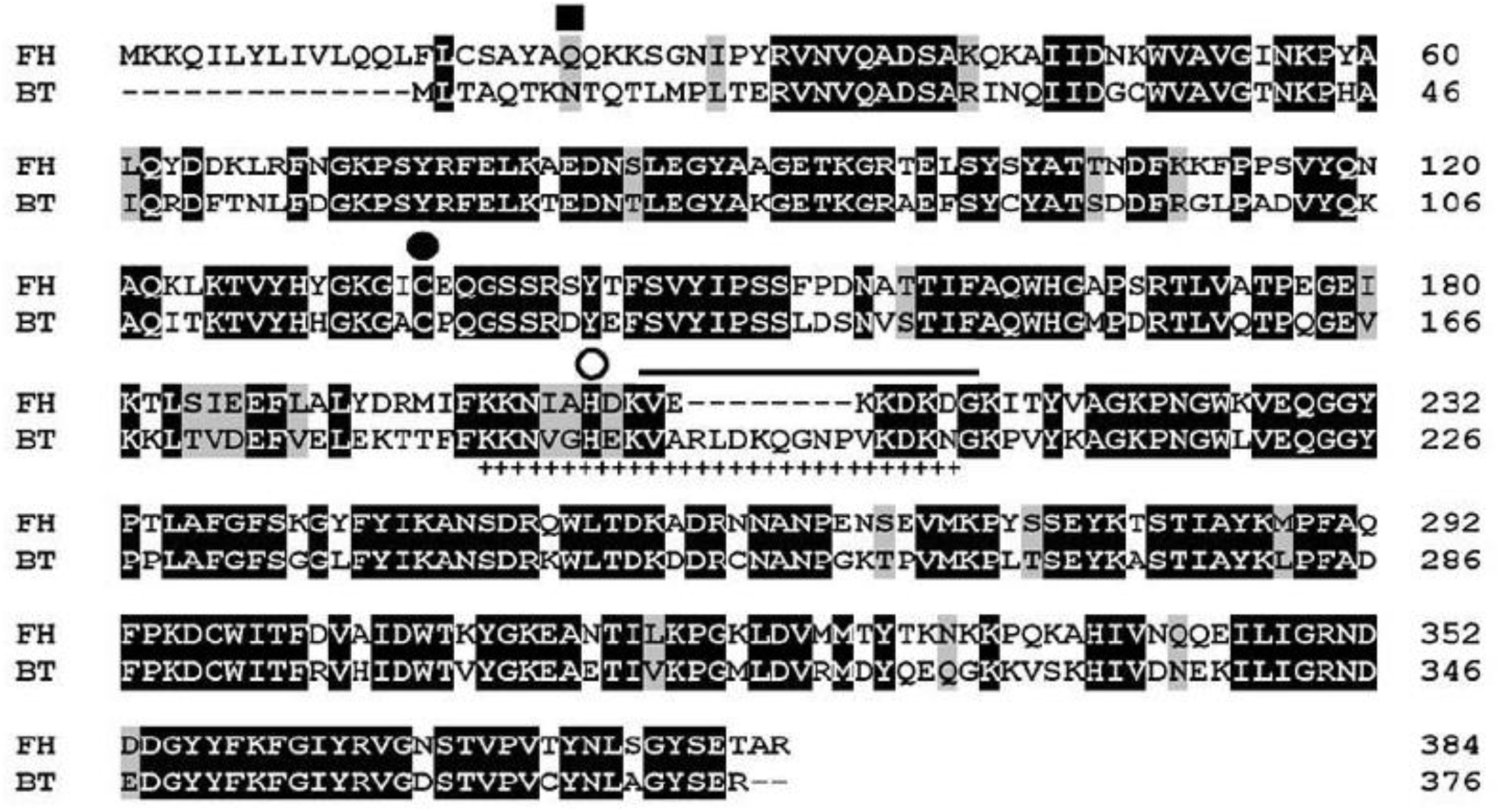
Alignment of the amino acid sequences of PhHepI and BtHepI.(Identical and similar residues are highlighted in black and gray, respectively. A closed rectangle indicates the putative signal peptide cleavage site at alanine-21 in F. heparinum heparinase I. Closed and open circles indicate conserved active site cysteine and histidine, respectively [26,27]. The solid overline indicates the putative calcium-coordinating domain [25]. The putative heparin-binding domain is indicated by the hatched underline. FH = F. heparinum; BT = B.thetaiotaomicron.) (Luo Y et.al. 2007)

On a base of homology between amino acid sequences of heparinases we found the areas closely to the active site for *Bacteroides thetaiotaomicron* HepI on its three-dimensional model structure and suggested the presence of similar areas for heparinase I *F. heparinum* (Fig. 2)

**Figure 2.**
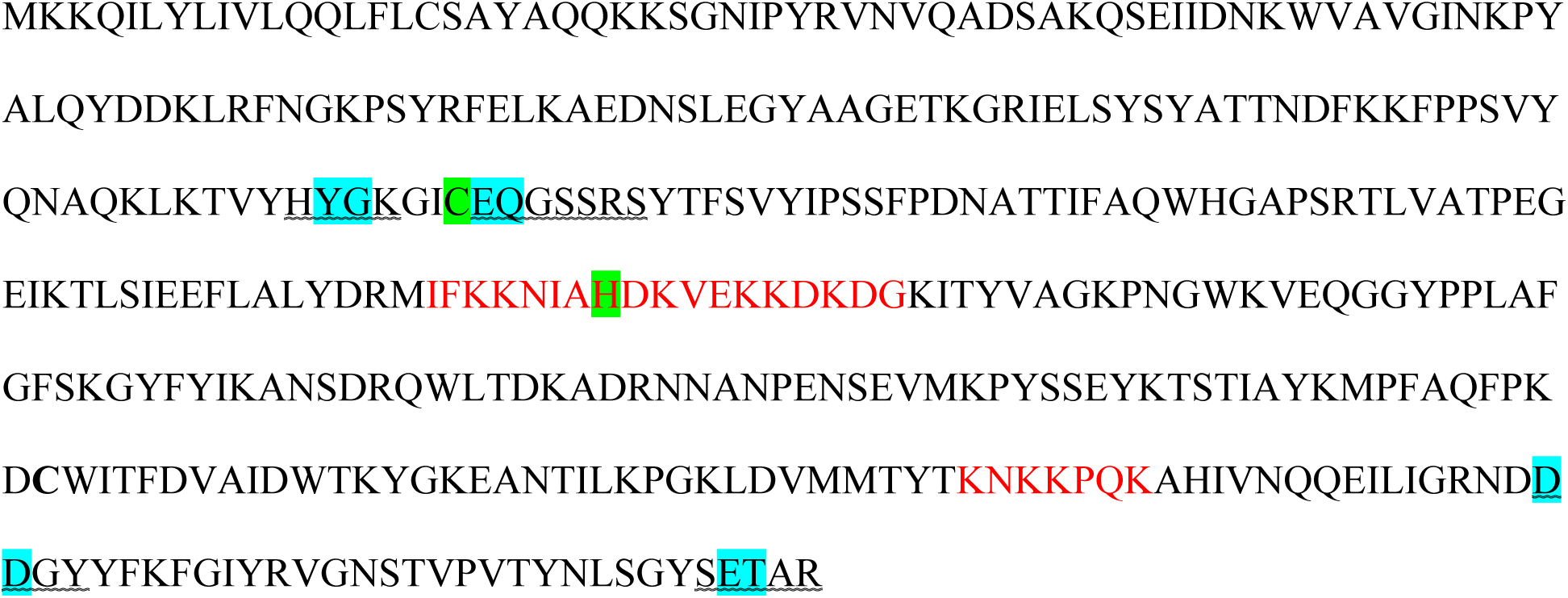
Heparinase I of Flavobacterium heparinum (sequence from GenBank) (C-135, H –203 – “Key” amino acids of the active center; IFK – substrate binding site; SET – supposedly, the area close to the active site of heparinase I (C-135); ET – amino acid residues for which mutations were introduced)

Mutations were introduced using degenerated primers, 2 amino acid substitutions for one act mutagenesis. To test the activity of mutant heparinase I vector pET22b (+) - Hep was constructed (Fig.3)

Screening of the obtained mutant heparinases was produced in compare with native heparinase I, by measuring the total activity heparinase I using the developed technique: Cells were grown as described in Materials and methods. Then pellet of cells was washed by distillated water and resuspended in 100**μl** lysozyme solution (3 mg/ml).The incubation was carried out at 37°C for 30 minutes.Then the cells were harvested by centrifugation (1min at 4 °C and 5000 rpm). The 50 μl supernatant was added to 25 μl heparin solution (17mM Tris–HCl, pH 7.0, 44mM NaCl, 3.5mM CaCl 2 and 25g/l heparin), mixed and incubated at at 30°C for 2h. The 50 μl of supernatant (after lysis cells) was warmed at 65°C for 15 min to inactivation of heparinase I, added to sane heparin solution, also incubated and used as a control sample. Twenty microliter of solution taken from the mixture were added to 2ml of 20mg/l Azure A to measure the increase in absorbance at 620nm.

The addition method was used for semi-quantitative analysis of the heparinase I activity: several dilutions of heparinase I F. heparinum (Sigma) were added to samle (supernatant after lysis of calles), in which heparinase I activity was necessary be measured. After that, measurement was performed by the method described above.

It should be noted that only two areas of the substitution from the four gave positive results. Combination of substitutions was constituted of the most effective mutation. So the best results were obtained with the introduction of the following substitutions E136 → Q, Q137 → P, E381 → P, T382 → P. For obtained mutant heparinase I activity was determined (Table 2, Fig. 4).

**Table 2.** The results of measuring the activity of the native heparinase I by semi-quantitative methods (data were averaged).

| Amount of added heparinase I, U | OD, 620nm | OD test sample – OD control sample |
| --- | --- | --- |
| 0 | 0,482 | 0,055 |
| 0,2 | 0,669 | 0,242 |
| 1 | 0,901 | 0,474 |
| 1,4 | 0,966 | 0,539 |
| 2 | 1,098 | 0,671 |
| control sample | 0,427 |  |

**Figure 3.**
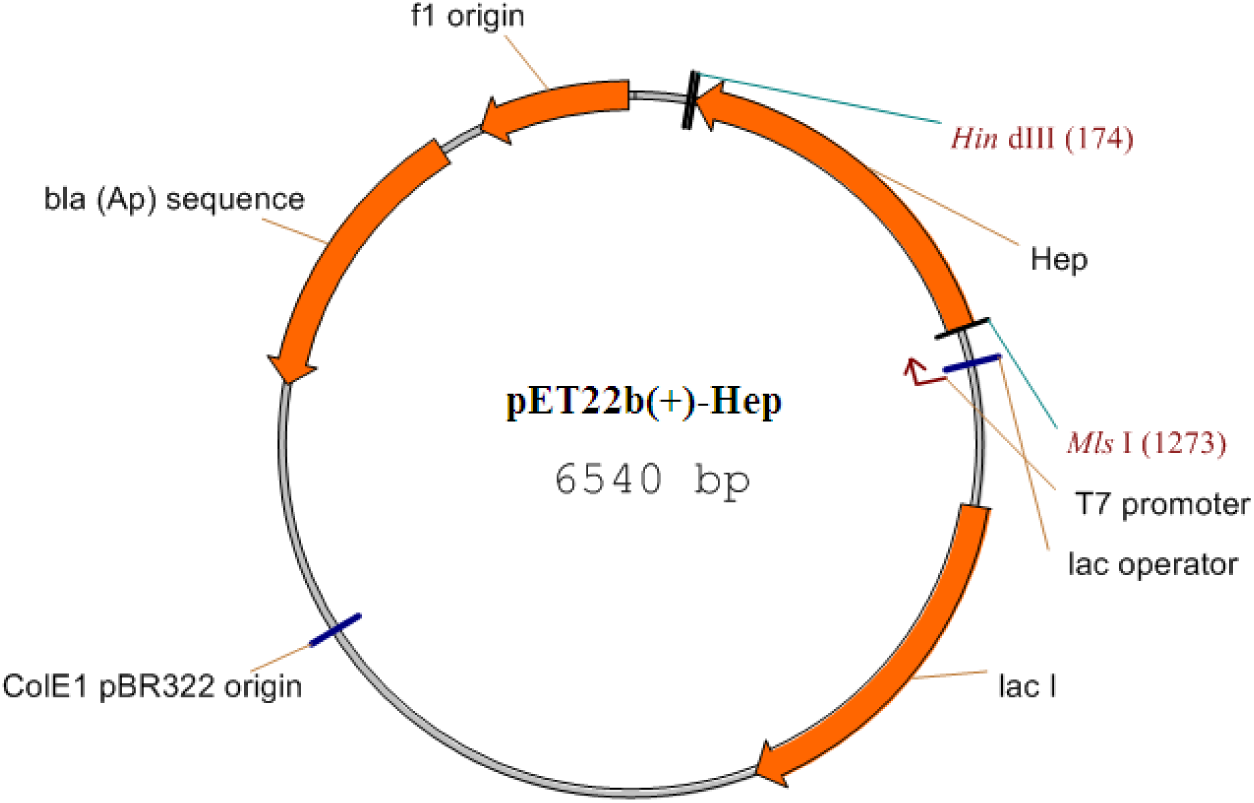
Costruction of expression vector pET22b (+) – Hep, that contain gene encoding heparinase I *F. heparinum* (Hep – gene encoding heparinase I, bla –gene encoding β-lactamase)

**Figure 4.**
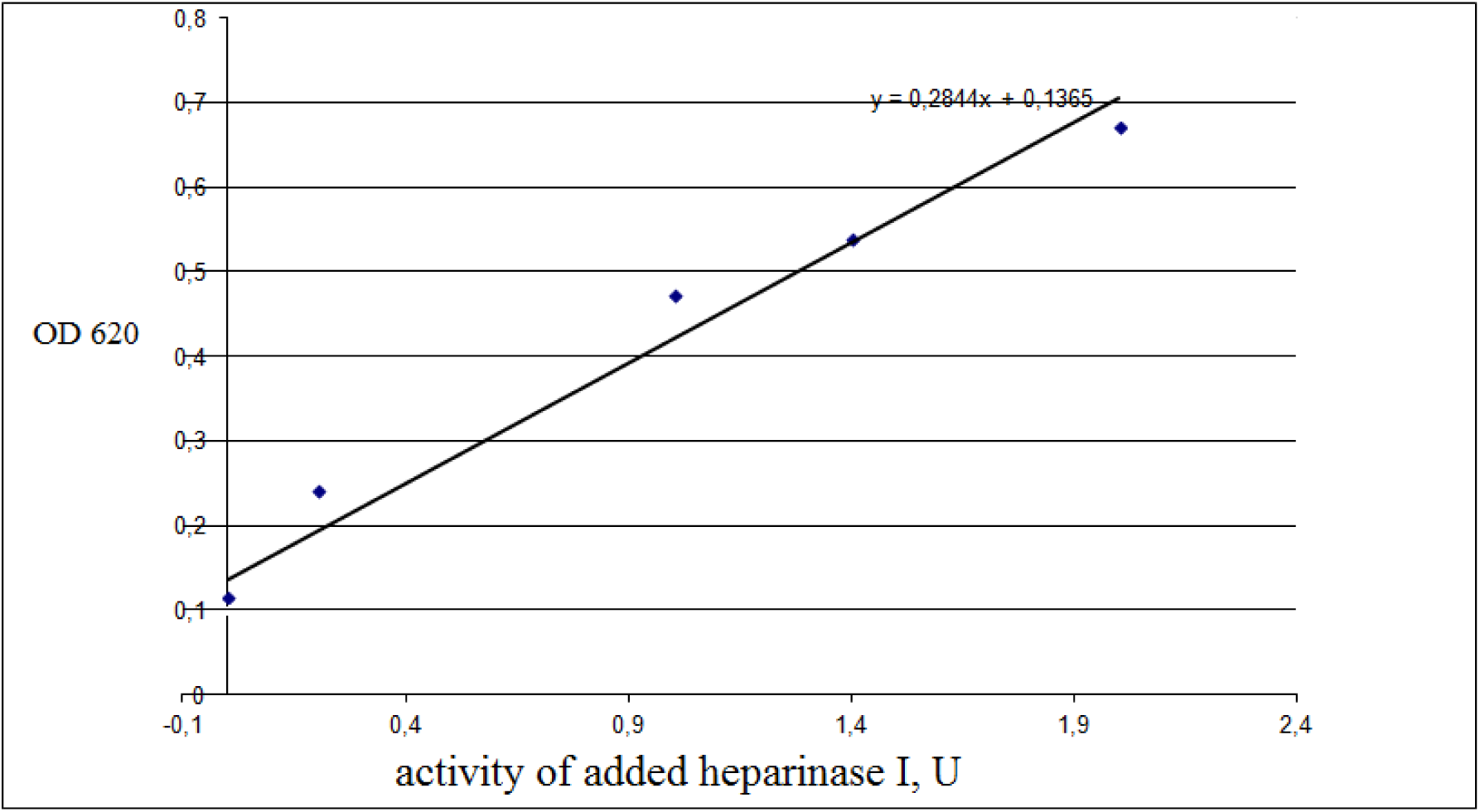
A graph of the change in absorbance solution of the Azur A control sample from amount of added heparinase I for the native heparinase I.

Also a graph of the change in absorbance solution of the Azur A was constructed for sample of mutant heparinase I. As a result the activity of the native heparinase I is 0.1365 units per sample, and the mutant heparinase I is 0,183 units, which is higher than the native.

Purifications of the mutant protein Hep-QP-PP and the native heparinase I were carried out and the specific heparinase I activities were measured. The results of purification of native and mutant heparinase I are shown in Fig. 5.

**Figure 5.**
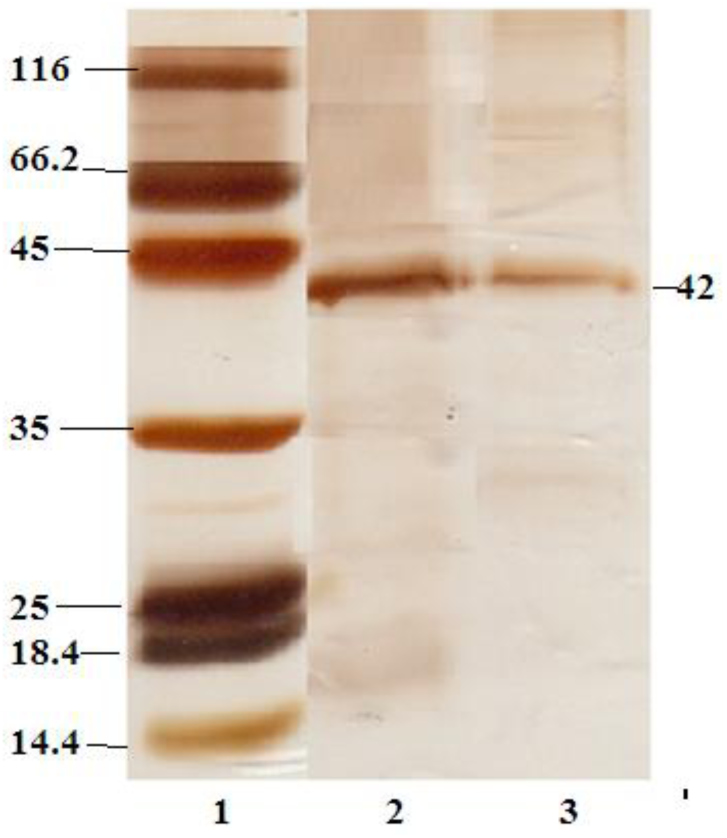
SDS–PAGE of proteins mutant and native heparinase I after purification (1-marker; 2 - mutant heparinase I; 3 - native heparinases I).

The determination of specific activities heparinase I was carried out by modification of the UV 232 nm method (Bernstein et al. 1988). Wherefore calibration graph of the change in absorption heparin solution for 1 hour at 232nm from the activity of heparinase I was constructed (Fig. 6).

**Figure 6.**
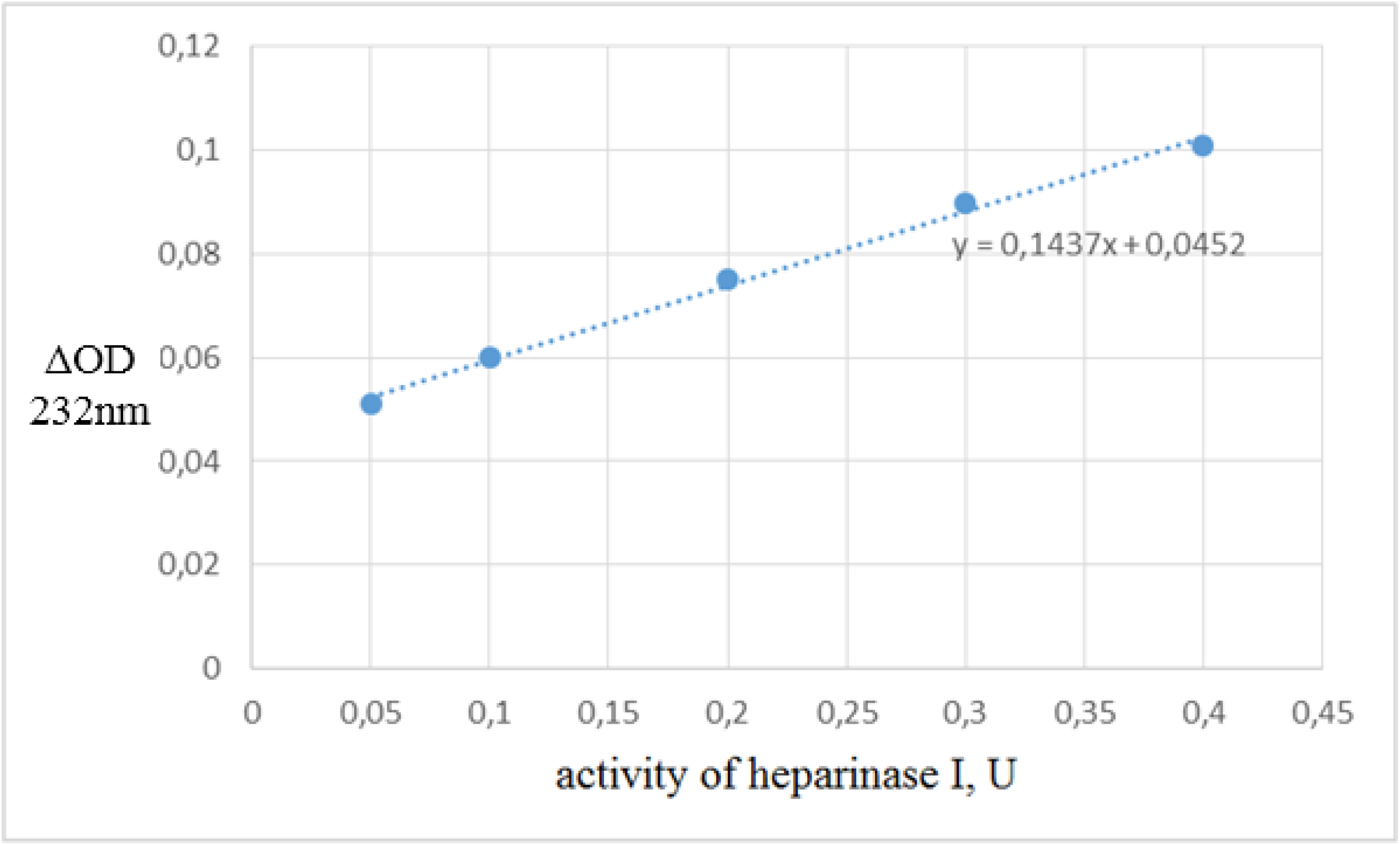
A calibration curve of the change in absorbance heparin solution for 1 hour at 232nm from amount of added heparinase I.

The table shows the results of measurements of the purified heparinase I (Table 3).

**Table 3.**
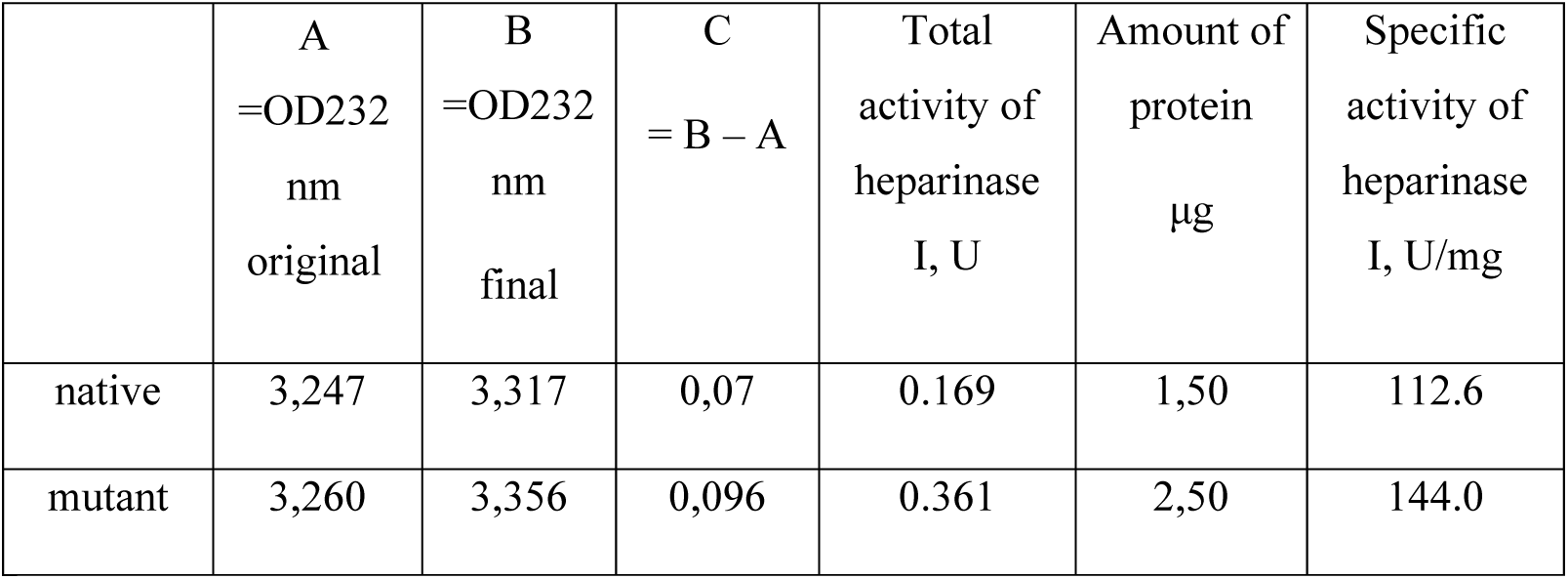
Results of measurement of the specific activity of heparinase I in samples.

The table shows that the specific activity of the native heparinase I is 112.6 U / mg, and the mutant heparinase I is 144.0 U / mg, which is about 25% higher than the native.

As a result of site-directed mutagenesis: the mutant heparinase I was obtained, comprising the following substitutions E136 → Q, Q137 → P, E381 → P, T382 → P, and having a specific activity on 25% (± 2%) higher than that of native.

## Acknowledgement

The work was supported by the Ministry of Education and Science of the Russian Federation (Project №075-15-2019-1658), and was performed using the resources of the National Bioresource Center – All-Russian Collection of Industrial Microorganisms, and the Center for Collective Use of GosNIIgenetik

